# Proteome specialization of anaerobic fungi during ruminal degradation of recalcitrant plant fiber

**DOI:** 10.1101/2020.01.16.907998

**Authors:** Live H. Hagen, Charles G. Brooke, Claire Shaw, Angela D. Norbeck, Hailan Piao, Magnus Ø. Arntzen, Heather Brewer, Alex Copeland, Nancy Isern, Anil Shukla, Simon Roux, Vincent Lombard, Bernard Henrissat, Michelle A. O’Malley, Igor V. Grigoriev, Susannah Tringe, Roderick Mackie, Ljiljana Pasa-Tolic, Phillip B. Pope, Matthias Hess

**Affiliations:** Faculty of Biotechnology, Chemistry and Food Science, Norwegian University of Life Sciences, Norway; University of California, Davis, CA, USA; DOE Environmental and Molecular Sciences Laboratory, Richland, WA, USA; Washington State University, Richland, WA, USA; U.S. Department of Energy Joint Genome Institute, Lawrence Berkeley National Laboratory, Berkeley, CA, USA; CNRS, UMR 7257, Université Aix-Marseille, 13288 Marseille, France; Institut National de la Recherche Agronomique, USC 1408 Architecture et Fonction des Macromolécules Biologiques, 13288 Marseille, France; Department of Biological Sciences, King Abdulaziz University, 21589 Jeddah, Saudi Arabia; Department of Chemical Engineering, University of California, Santa Barbara, CA, USA; Department of Plant and Microbial Biology, University of California, Berkeley, CA, USA; Department of Animal Science, University of Illinois, Urbana-Champaign, IL, USA; Faculty of Biosciences, Norwegian University of Life Sciences, Norway

**Author notes:** Corresponding author: Live H. Hagen –.

## Abstract

The rumen harbors a complex microbial mixture of archaea, bacteria, protozoa and fungi that efficiently breakdown plant biomass and its complex dietary carbohydrates into soluble sugars that can be fermented and subsequently converted into metabolites and nutrients utilized by the host animal. While rumen bacterial populations have been well documented, only a fraction of the rumen eukarya are taxonomically and functionally characterized, despite the recognition that they contribute to the cellulolytic phenotype of the rumen microbiota. To investigate how anaerobic fungi actively engage in digestion of recalcitrant fiber that is resistant to degradation, we resolved genome-centric metaproteome and metatranscriptome datasets generated from switchgrass samples incubated for 48 hours in nylon bags within the rumen of cannulated dairy cows. Across a gene catalogue covering anaerobic rumen bacteria, fungi and viruses, a significant portion of the detected proteins originated from fungal populations. Intriguingly, the carbohydrate-active enzyme (CAZyme) profile suggested a domain-specific functional specialization, with bacterial populations primarily engaged in the degradation of polysaccharides such as hemicellulose, whereas fungi were inferred to target recalcitrant cellulose structures via the detection of a number of endo- and exo-acting enzymes belonging to the glycoside hydrolase (GH) family 5, 6, 8 and 48. Notably, members of the GH48 family were amongst the highest abundant CAZymes and detected representatives from this family also included dockerin domains that are associated with fungal cellulosomes. A eukaryote-selected metatranscriptome further reinforced the contribution of uncultured fungi in the ruminal degradation of recalcitrant fibers. These findings elucidate the intricate networks of *in situ* recalcitrant fiber deconstruction, and importantly, suggests that the anaerobic rumen fungi contribute a specific set of CAZymes that complement the enzyme repertoire provided by the specialized plant cell wall degrading rumen bacteria.

## Introduction

It has been estimated that there are approximately 1 billion domesticated ruminant animals^1^ and numbers are predicted to increase further in order to provide food security for the growing human population^2^. The societal importance of ruminants has fueled global efforts to improve rumen function, which influences both animal health and nutrition. In particular, broadening the knowledge of the complex microbial interactions and the enzymatic machineries that are employed within the rumen microbiome is thought to provide means to efficiently optimize feed conversion, and ultimately the productivity and well-being of the host animal.

One of the major functions mediated by the rumen microbiome is to catalyze the breakdown of plant carbon into short-chain fatty acids (SCFA) that can be metabolized by the host animal. To facilitate the degradation of complex plant carbohydrates, the rumen microbiome encodes a rich repertoire of carbohydrate-active enzymes (CAZymes). This group of enzymes is categorized further into different classes and families, which include carbohydrate-binding modules (CBMs), carbohydrate esterases (CEs), glycoside hydrolases (GHs), glycosyltransferases (GTs), and polysaccharide lyases (PLs)^3^. Previous studies have mostly been dedicated to CAZymes from rumen bacteria, although it is becoming increasingly clear that fungi and viruses also possess key roles in the carbon turnover within the rumen^4,5^. Over the last decade, targeted efforts to isolation and cultivate novel rumen microorganisms have resulted in a more detailed understanding of the physiology of anaerobic rumen archaea and bacteria and their contribution to the overall function of the rumen ecosystem^6^. Recent studies have also shed light on the viral rumen population and although work in this area is still nascent, it suggests that the rumen virome modulates carbon cycling within the rumen ecosystem through cell lysis or re-programming of the metabolism of the host microbiome^5,7,8^. Anaerobic rumen ciliate protozoa and fungi have largely remained recalcitrant to both cultivation and molecular exploration efforts^9^, and although recent cultivation efforts have provided important insight into the lifestyle and enzymatic capacity^4,10^, their quantitative metabolic contributions to the greater rumen ecosystem are still unclear.

Enumerating anaerobic rumen fungi is challenging, mainly due to their different life stages and their growth within plant fragments as well as sub-optimal DNA extraction and molecular methods to recover their genomic information^11–13^. Reported counts of fungal cells vary greatly between studies, with numbers ranging between 10^3^ and 10^6^ cells/ml of rumen fluid^14–16^. To date, only a total of eighteen genera (*Agriosomyces, Aklioshbomyces, Anaeromyces, Buwchfawromyces, Caecomyces, Capellomyces, Ghazallomyces, Cyllamyces, Feramyces, Joblinomyces, Khoyollomyces, Neocallimastix, Liebetanzomyces, Oontomyces, Orpinomyces, Pecoramyces, Piromyces*, and *Tahromyces*), all belonging to the early-branching phylum Neocallimastigomycota, have been described^4,17–19^, although culture independent studies have suggested that this only represents half of the anaerobic fungal population that exist in the rumen ecosystem^17,20^. Genomes obtained from representatives of this phylum have been recognized to encode a large number of biomass-degrading enzymes and it is becoming increasingly clear that these currently still understudied organisms play a key role in the anaerobic digestion of complex plant carbohydrates^4,10,21^. The impact of fungi in the rumen ecosystem was already demonstrated in the early 1990s by Gordon and Phillips who reported a significant decrease in fiber digestion within the rumen after anaerobic fungi had been removed by the administration of fungicides^22^. The importance of rumen fungi for biomass degradation has since then been supported by *in vivo* studies^23–25^, and recently reinforced in transcriptome studies revealing that the fungi express a range of CAZymes when grown on different carbon sources^9,26^. Although enzymes of fungal origin have been regularly explored for their remarkable capacity to degrade lignocellulosic fiber^12,27,28^, their functional role in native anaerobic habitats and within the biomass-degrading enzyme repertoire of the rumen microbiome remains unclear. Thus, we lack a complete understanding of their biology and their contribution to the function and health of the rumen ecosystems.

To fill this knowledge gap, we utilized a genome-centric metaproteome approach to investigate the distinct role of the fungal population during the biomass-degradation process in the rumen. Moreover, our experiments were designed to target populations actively degrading recalcitrant fibers that resisted initial stages of microbial colonization and digestion. Specifically, metaproteomic data were interrogated using a database constructed from five available rumen fungal isolates^4^ in addition to genomes and metagenome-assembled genomes (MAGs) of cultured and uncultured rumen bacteria, respectively. To further explore the activity of uncultured fungi, we performed a second metaproteomic search against a database generated from polyadenylated mRNA extracted from rumen-incubated switchgrass. Combining data from these various layers of the rumen microbiome enabled us to generate new insights into the functional role of anaerobic rumen fungi, expanding our holistic understanding of plant-fiber decomposition in the rumen ecosystem.

## Results & Discussion

### Taxonomic origin of proteins involved in rumen biomass-degradation

To directly link the microbial genes actively involved in the degradation of complex plant material in the anaerobic rumen ecosystem, we incubated milled switchgrass in *in situ* nylon bags within the rumen of two cannulated dairy cows to encourage colonization by the native rumen microbiota. After an incubation period of 48 hours, bags were collected, and proteins were extracted from the rumen-incubated fiber for metaproteome profiling. To resolve the roles of the fungal, bacterial and viral populations, we designed a customized <u>RU</u>men-<u>S</u>pecific <u>ref</u>erence <u>D</u>ata<u>B</u>ase (hereby referred to as ‘RUS-refDB’). To specifically determine the metabolic function of the fungal population, the genomes of five rumen fungi that were available at the time of our data analysis [i.e. *Anaeromyces robustus, Neocallimastix californiae, Pecoramyces ruminantium* C1A (formerly classified as *Orpinomyces* sp. C1A), *Piromyces finnis*, and *Piromyces* sp. E2^4,18,21,29^] were included in the database. The RUS-refDB was further complemented with 103 metagenome-assembled genomes (MAGs) and 913 metagenome-assembled viral scaffolds (MAVS) recovered from a rumen metagenome we generated previously using a comparable experimental design of rumen-incubated switchgrass^30,31^. To ensure that the database also represented the major functional and phylogenetic groups of well-known rumen prokaryotes, we searched the Hungate1000 collection^6^ and selected the genomes of 11 cultured rumen bacteria, including species related to *Ruminococcus, Prevotella* and *Butyrivibrio*. We also included the genomes of *Fibrobacter succinogenes* S85^32^ and *Methanobrevibacter ruminantium* M1^33^, both shown to play a significant role in proper rumen function. A summary of the MAGs, MAVS and isolated genomes contributing to our custom-built RUS-refDB is provided in **Supplementary Table S1.** Mapping the protein scans from switchgrass fiber and rumen fluid against the RUS-refDB resulted in the identification of a total of 4,673 protein groups, and a strong positive correlation (Pearson correlation R > 0.8) of the two biological replicates (cow 1 & cow 2) was obtained (**Supplementary Figure S1**).

To obtain an overview of the microbial taxa associated with our detected proteins, we generated a phylogenetic tree and included the numerical detection of proteins for each taxon (**Figure 1**, numerical detection of proteins can be found in **Supplementary Table S1**) in both the switchgrass fiber fraction and rumen fluid. Interestingly, the (meta)genome-resolved metaproteome revealed that a high fraction of detected proteins within our metaproteome were of fungal origin. Within the five anaerobic rumen fungi, we observed between 316 and 787 proteins that aligned well to proteins predicted from the genomes of *Piromyces finnis* and *Neocallimastix californiae*, respectively. This exceeds the number of proteins detected from any of the investigated prokaryotes included in this study, and likely reflects the fundamental functional role that fungi hold in ruminants during degradation of recalcitrant cellulosic material. Moreover, the metaproteomics data also revealed a higher level of protein grouping across the fungal genomes due to homologues proteins, suggesting that there are conserved features of the fungal genomes that have been sequenced to date. Many of the corresponding protein-coding genes were also replicated within each fungal genome, demonstrating that individual rumen fungi hold several sets of functionally important genes. Despite a reportedly high degree of horizontal gene transfer (HGT) in the rumen microbiome^4,34,35^, only a few detected proteins mapped to both fungi and prokaryotes, suggesting that the overall sequences of these particular enzymes are evolutionary divergent across these two kingdoms.

**Figure 1:**
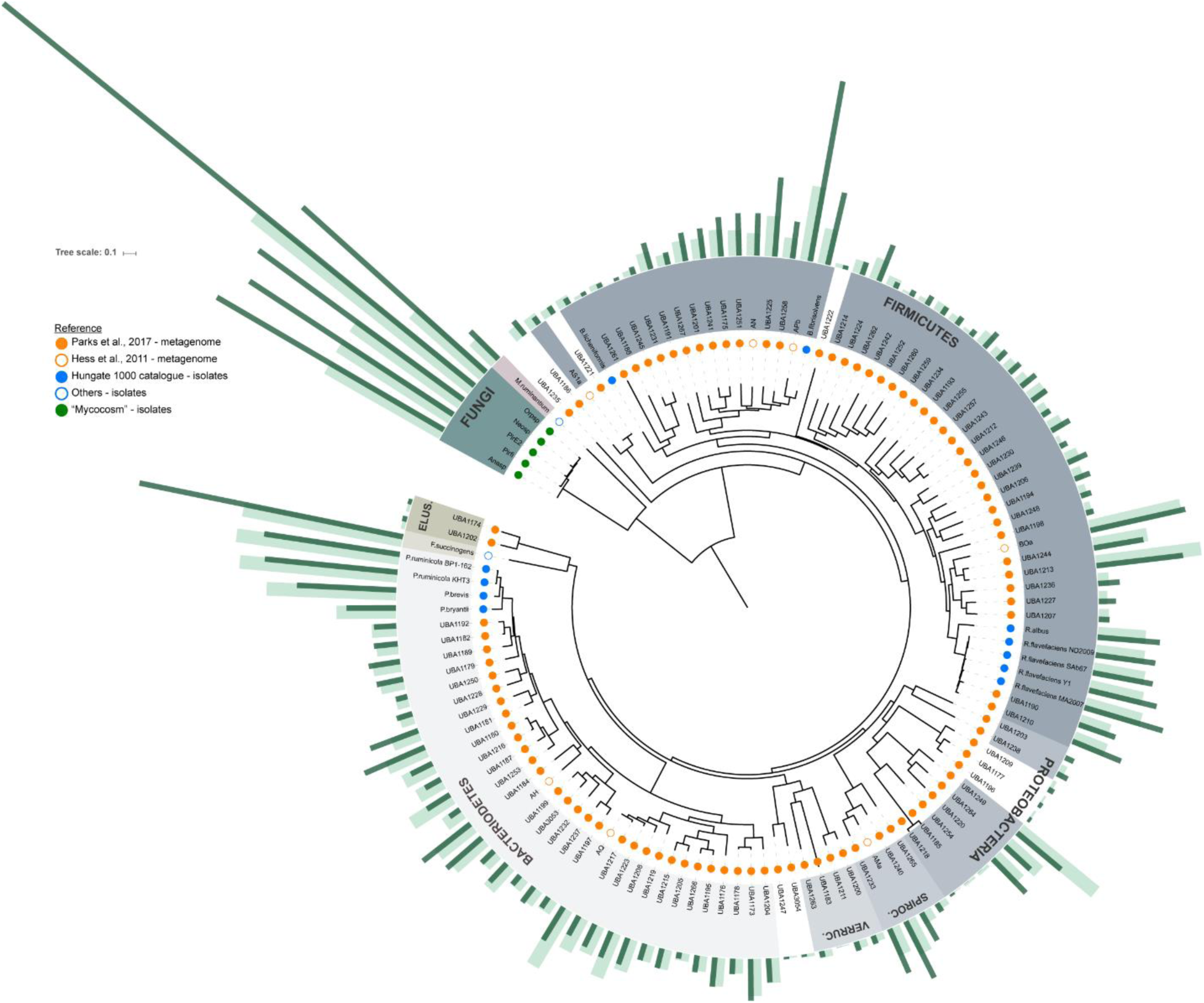
Concatenated ribosomal protein tree of the genomes and metagenome-assembled genomes (MAGs) included in RUS-refDB. Phyla-level groups are colored in shades of grey (bacteria), magenta (archaea) and green (fungi: Anasp, *Anaeromyces robustus;* Pirfi, *Piromyces finnis; PirE2, Piromyces* sp. E2; Neosp, *Neocallimastix californiae*; Orpsp; *Orpinomyces* sp.), and labeled inside the circle (Spiroc., Spirochaetes; Verruc., Verrucomicrobia; Elus., Elusimicrobia). MAGs/clades with uncertain taxa have white background. Circles at the end of each node are color coded by the metagenome data set or genome collection each MAG/genome in RUS-refDB originated from, as indicated in the top left legend. The number of detected proteins from the samples in the switchgrass fiber fraction (dark green) and rumen fluid (light green) are specified by bars surrounding the tree. In cases where a protein group consisted of two or more homologues protein identifications, each protein match is considered. The viral scaffolds, not included in the tree, had 56 and 62 proteins detected in switchgrass fiber and rumen fluid respectively. Numerical protein detection can be found in **Table S1**. A complete version of this tree is available in Newick format as **Supplementary Data S3.**

The bacterial portion of the RUS-refDB was mostly comprised of genomes belonging to the Firmicutes and Bacteroidetes phyla, of which species belonging to the *Ruminococcus* and *Prevotella* accounted for a large fraction of the detectable proteins (**Figure 1**). A high number of detected proteins also aligned well to the genome of *Butyrivibrio*, emphasizing the significance of this group in biomass-degradation and conversion within the rumen. Within this clade, ‘APb’, a MAG of an as-yet uncultured prokaryote, phylogenetically closely related to *Butyrivibrio fibrisolvens*, showed the highest number of detected proteins (switchgrass: 237; rumen fluid: 97). Not unexpectedly, the well-studied fibrolytic bacteria *Fibrobacter succinogenes* represented the bacterial species with the highest number of detected proteins (switchgrass: 349; rumen fluid: 210), followed by two strains of *Prevotella ruminicola* (ranging from 173 to 213 proteins, of which the majority of the detection proteins were homologues of the two strains) and *P. brevis* (switchgrass: 129; rumen fluid: 168), highlighting their overall importance in the carbohydrate metabolisms in the rumen. This is consistent with previous studies involving functional analysis of the rumen microbiome, demonstrating that a majority of the plant cell wall polysaccharide degradation is carried out by species related to *Fibrobacter, Ruminococcus* and *Prevotella*^24,25,36^. Although our metaproteome data suggested that these aforementioned characterized prokaryotes were amongst the most active (i.e. highest numbers of detected proteins), a significant fraction of the protein groups mapped to MAGs representing uncultured and uncharacterized taxa. This included MAGs classified within the *Bacteroidetes* phyla, such as UBA1181 previously described by Naas et al.^37^, a clade consisting of the *Spirochaetes*-assigned MAG ‘AMa’, UBA1233 and UBA1240, in additional to a *Proteobacteria*-clade (UBA1249, UBA1220 and UBA1264). This reiterates that a considerable fraction of the bacterial rumen microbiome remains to be explored and characterized before a holistic and truly advanced understanding of the role of rumen bacteria is achieved.

### Metaproteome-generated CAZyme profile indicates compartmentalized niches amongst fungal and bacterial populations

The efficiency of the rumen microbiome in breaking down the complex cell wall of plants is due to the orchestrated synthesis, degradation, and modification of glycosidic bonds by an intricate mixture of microorganisms and their CAZymes. Crystalline cellulose is often degraded through a synergistic mechanism between endo- and exo-acting CAZymes targeting the glycosidic bonds within or at the ends of the polysaccharide, respectively. To visualize the specific enzyme-contrived contributions of the different microbial taxa during plant biomass digestion within the rumen ecosystem, we analyzed and constructed CAZy profiles of the detected proteins from each predicted source organism (**Figure 2**).

**Figure 2:**
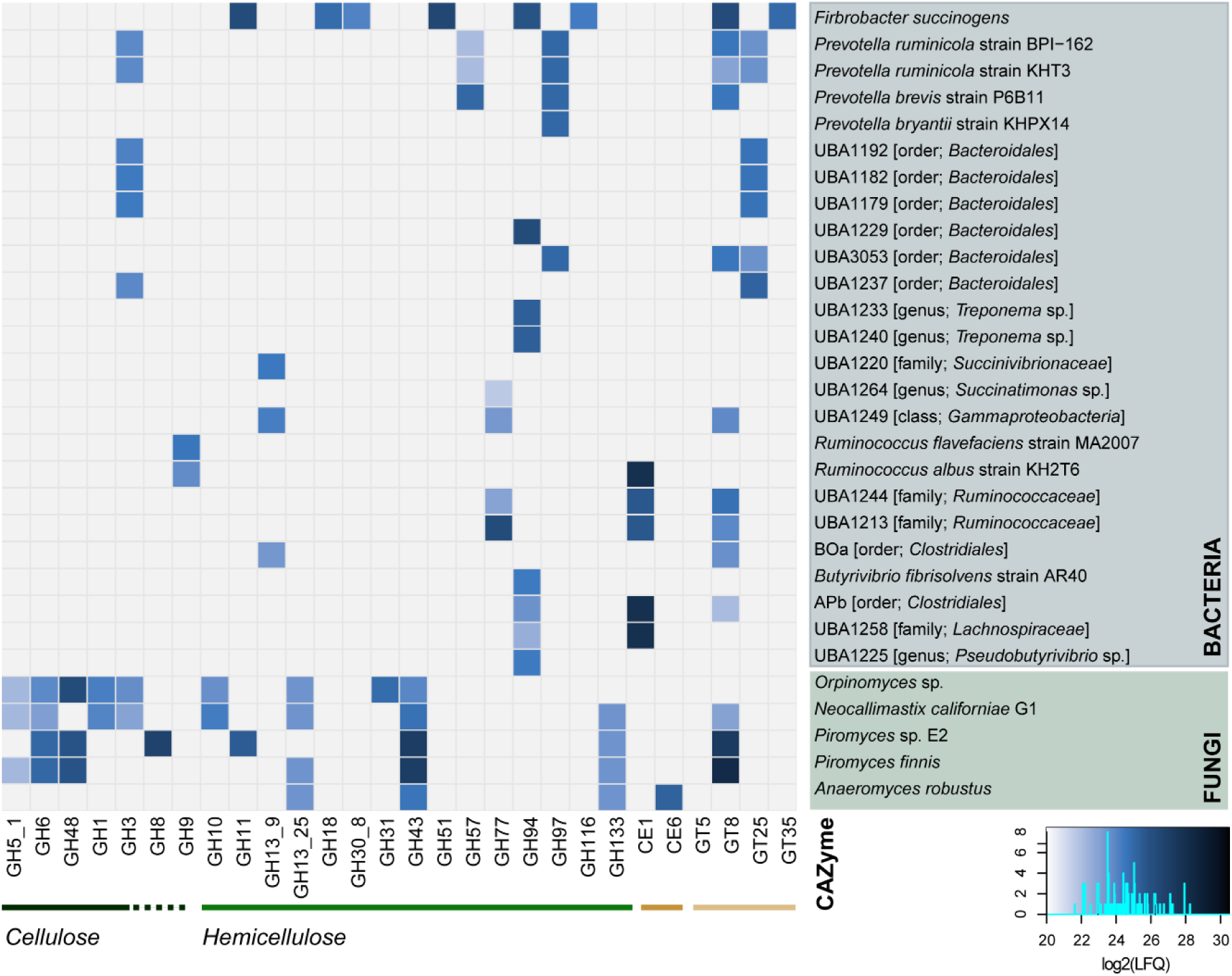
CAZyme profile from each predicted source organism in RUS-refDB, displaying the detected proteins associated with the milled switchgrass. Here, we focused only on CAZymes detected in both animals to achieve high confidence detection of the active key populations. The colors in the heat map indicates the protein detection levels of each protein group reported as the average Log2(LFQ)-scores for the biological replicates, where a light blue color is low detection while darker is high protein detection.

Interestingly, the exo-acting cellulases with highest protein abundance (measured by Label-Free Quantification (LFQ)) in our dataset, such as GH6 and GH48, appeared to come nearly exclusively from the rumen fungi. Moreover, GH48, aligning well to predicted protein sequences from both *Orpinomyces* sp. and the two *Piromyces* species, had the highest LFQ-level of all cellulases. While GH48 have been absent or only detected at very low levels in previous rumen metagenomes^31,38^, members of this GH family have been observed in rumen metatranscriptomes from mixed rumen populations, reportingly expressed by *Ruminococcus* and rumen fungi^24,25,39^. It is also worth noting that while members of the GH48 family were the most abundant CAZymes affiliated to protein sequences of *Orpinomyces* sp. origin, other detected proteins belonging to the glycoside hydrolase families GH1, GH3, GH5_1, and GH6 also aligned to the proteome of this fungus. In contrast to the elevated number of mapped proteins (**Figure 1**), CAZymes predicted from the genome of *N. californiae* contributed at lower detection levels than its fungal companions, albeit its CAZyme profile covered proteins with a wide range of substrate specificity including both cellulose (i.e. β-glucosidases, GH1 and GH3; endoglucanases, GH5_1 and GH6), starch (amylase and amyloglucosidases, GH13_25-GH133) and hemicellulose (xylanases, GH10 and GH43). CAZymes inferred in the conversion of starch and hemicellulose also aligned well to the four other fungal reference genomes, with elevated level of xylanases belonging to the GH43 family (**Figure 2, Supplementary Data S1**).

Despite the cellulose-degrading reputation of *F. succinogenes*, the detected CAZymes were predominately involved in soluble glucans and/or hemicellulose degradation (**Figure 2**), with representatives belonging to the family of GH11, GH51 and GH94 amongst the most abundant glycoside hydrolases. In addition to *F. succinogenes, R. albus* and *R. flavefaciens* have also been repeatedly shown to contribute many of the required CAZymes for biomass-degradation in the rumen^36,40–42^. Indeed, endoglucanase GH9, a CAZyme family capable of hydrolyzing β 1→ 4 glyosidic bonds in cellulose, were detected in the proteome of both *R. albus* and *R. flavefaciens*. Members of the previously mentioned GH48-family, that suggested *Ruminococcus* sp. as key to cellulose degradation^43,44^, however, were only detected at low confidence levels (i.e. not found in replicates) and at very low LFQ levels (**Supplementary Data S1**). While our metaproteome data confirmed the enzymatic machineries of the previously mentioned characterized bacteria, proteins associated with recalcitrant cellulose decomposition were not restricted to these. The MAG of the uncultivated UBA1213, classified as a member of *Ruminococcaceae*, was associated with multi-domain proteins containing GH77 and GT35 at high abundance, whereas a close relative of UBA1213, ‘BOa’, mapped to multi-domain CAZymes possessing an α-amylase domain (i.e. GH13_9) and the carbohydrate binding module CBM48. Both these modules have been shown to be involved in starch degradation^45,46^, and our metaproteome further suggested that these two MAGs also expressed several enzymes involved in fermentation of starch-derived sugars (i.e. glycolysis, **Figure 4**). It should also be noted that a higher number of proteins aligned to those predicted for both UBA1213 and BOa compared to their cultivated *Ruminococcus* relatives (**Figure 1**). Besides *F. succinogenes* and *Butyvibrio fibrisolvens*, several MAGs (i.e. UBA1229, UBA1233, UBA1240 at high levels and ‘APb’, UBA1225 and UBA1258 at lower levels) also displayed significant protein detection levels of GH94, suggesting that cellobiose phosphorylation mediated through the action of GH94 is widespread amongst the rumen microbiome. Overall, it appears that within our experimental constraints (switchgrass incubated for 48 hours), bacterial populations contributed CAZymes that primarily modified non-cellulosic plant carbohydrates. It should be emphasized that the metaproteome data analyzed here represents only a snapshot of the community metabolism, and that the protein profiles of different rumen populations most likely have undergone temporal transformations in the time period between the plant material being introduced into the rumen environment and our analysis^23,47^.

**Figure 3:**
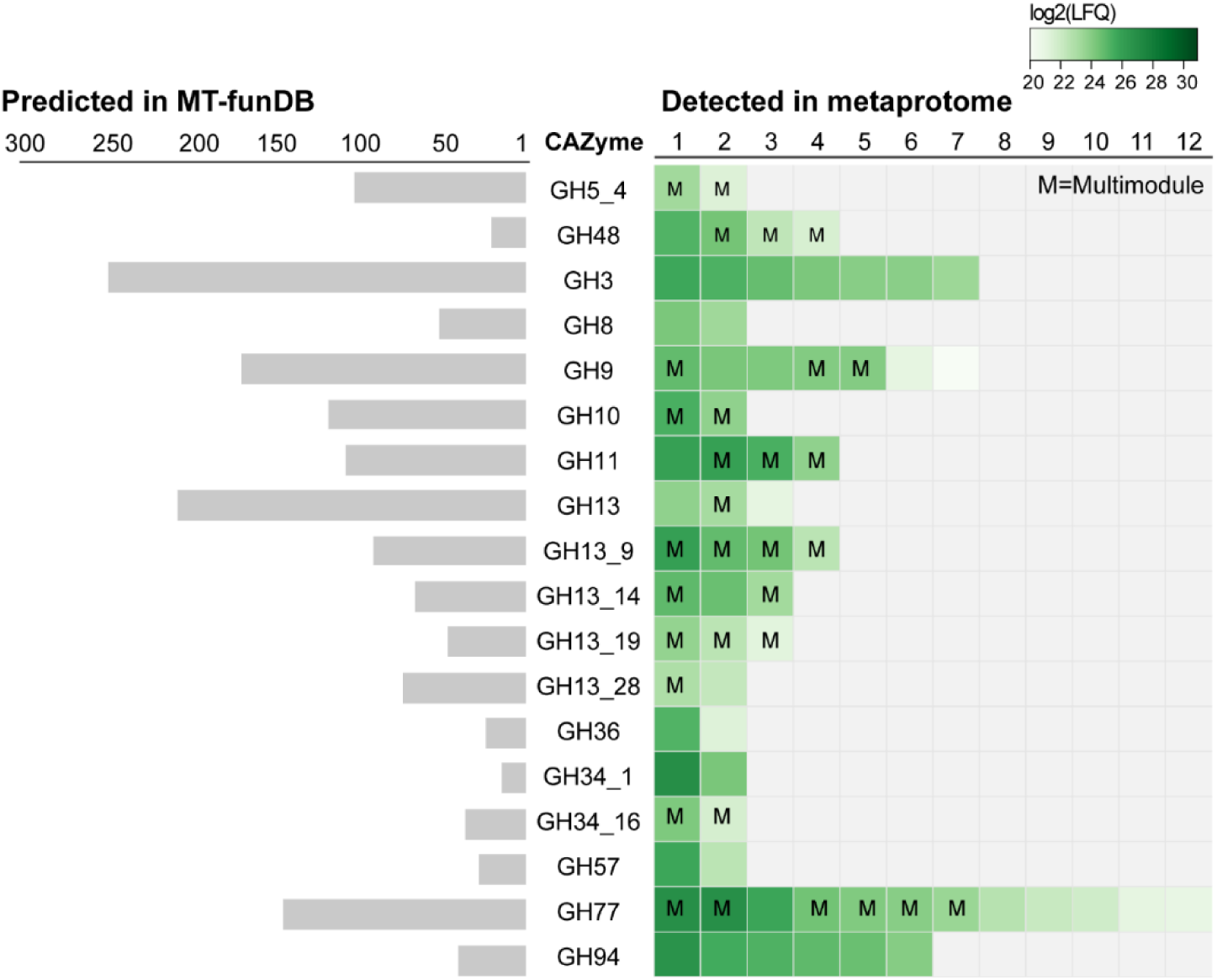
Visualization of the number of predicted genes annotated to specific GH families in MT-funDB (left) and those detected when searching MT-funDB against the metaproteome (right). Only CAZymes detected in both animals in at least one of the microhabitats are included to achieve high confidence detection. The colors of the squares in the left panel indicated the protein detection level for each individual protein, reported as the average log2(LFQ) of the biological replicates, where light green represents low detection level while darker green is high protein detection level. While this figure only shows those detected in the milled switchgrass, a comprehensive table of all CAZymes detected in both switchgrass and rumen fluid can be found in **Supplementary Data S2**. This also includes details regarding proteins with multiple CAZyme modules (indicated with an ‘M’).

### High prevalence of multi-modular domains and cellulosomal proteins

Some of the most efficient biomass degrading anaerobes possess cellulosomes, which are multienzyme complexes that enable the orchestrated and synchronized activity of various enzymes that are needed to degrade the cellulosic and hemicellulosic components of recalcitrant plant material^48^. Until recently, cellulosomes and their essential building blocks have been identified and described only from anaerobic bacteria^40,49–51^. However, advances in the isolation and cultivation of anaerobic fungi coupled with genome and transcriptome analyses have confirmed the presence of cellulosomes in anaerobic fungi for the well-synchronized deconstruction of plant carbohydrates^4^. Many CAZymes appear in multidomain modules, often comprising substrate-binding domains in addition to one or several domains specific for multifunctional GH families. Within our ruminal metaproteome, we detected proteins containing cellulosomal domains such as bacterial and fungal dockerins and carbohydrate-binding modules, which are specific for these large, multiprotein structures. These non-catalytic domains have recently been demonstrated to be numerous in anaerobic fungi, with an average of more than 300 non-catalytic dockerin domains encoded in the genome of each strain^4^. Accordingly, a significant number of the detected CAZymes in our metaproteome data contained at least one dockerin domain, with a clear preponderance of dockerins of fungal origin (**Table 1, Supplementary Table S2**). In general, while the bacterial cellulosome signature sequences encompassed a single Type-I dockerin (DOC1), the fungal counterparts frequently occurred as double or triple dockerins domains (here classified as type-II; DOC2). Dockerin domains in tandem repeats are indeed associated with fungal cellulosomes, and it is believed that this construction facilitates the involvement of more binding sites, thus binding potential substrates more efficiently, than single dockerins^4,52^.

**Table 1:**
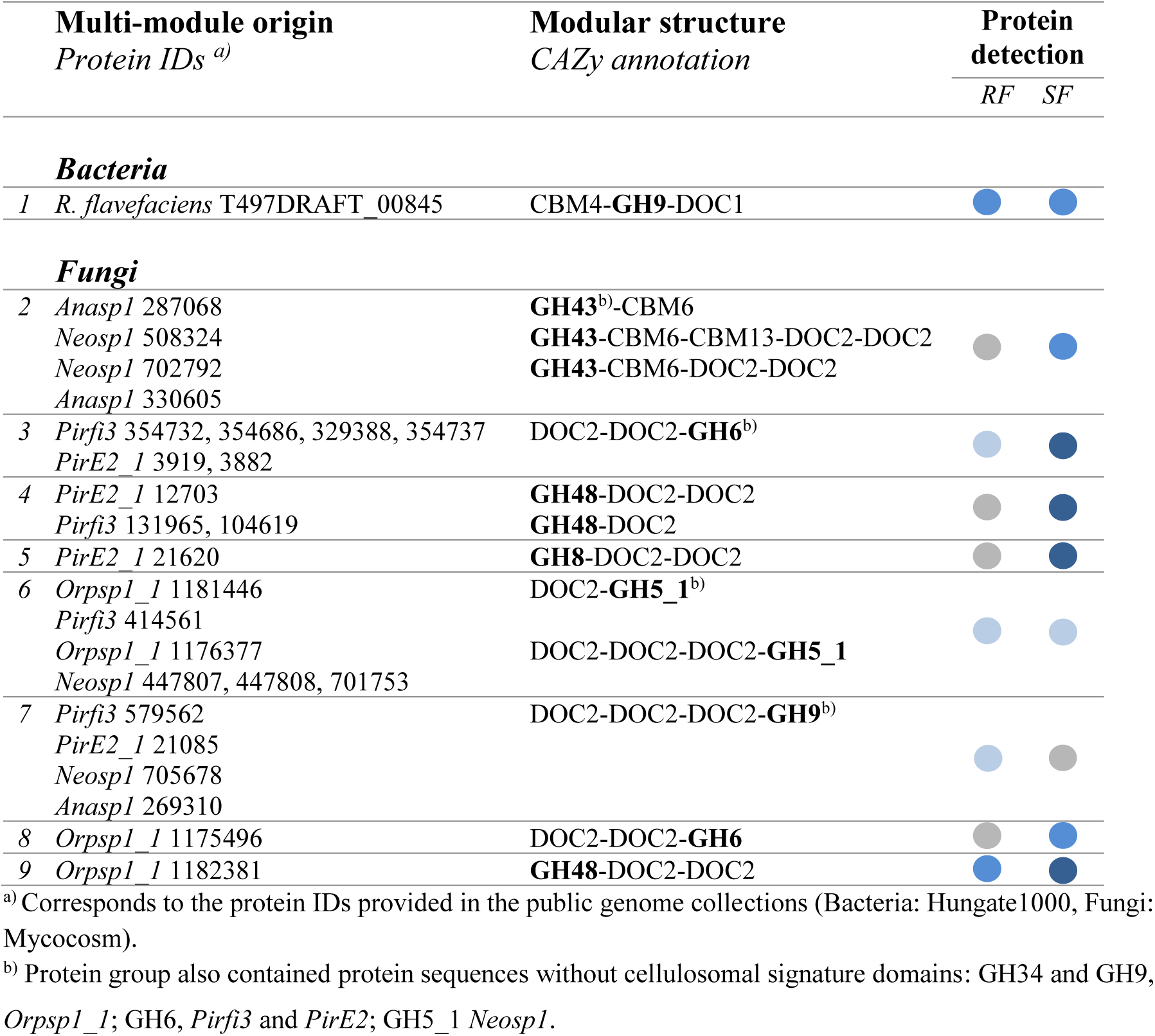
Cellulosomal subunits expressed during degradation of lignocellulosic biomass. Protein detection (LFQ) levels are indicated with color coded circles (light blue is low protein detection while dark blue is high. Grey is absent/below detection level), given as the average of the two animals in samples from rumen fluid (RF) and switchgrass fiber (SF). This table only contains protein groups detected in both animals in at least one of the microhabitats (SF and/or RF). An extended table is provided in **Supplementary Table S2**. Proteins of which only shared peptides was detected are grouped (i.e. protein groups) and quantified together. Nevertheless, these may have divergent domains within the sequence, that are not detected in our metaproteome.

The CAZymes containing dockerin domains in tandem repeats were further flanked with a variety of glycoside hydrolase domains, including those belonging to the GH3, GH5_1, GH6, GH8, GH9, GH43 and GH48 family. Notably, while GH3 and GH6 have recently been confirmed in fungal cellulosomes^4^, they seem to be absent in bacterial counterparts. Moreover, the GH48 enzymes detected in our metaproteome, except those affiliated with *Piromyces finnis*, contained two copies of dockerin domains (i.e. *Orpinomyces* sp., and *Piromyces* sp. E2, **Table 1**; *Anaeromyces robustus* and *Neocallimastix californiae*, **Supplementary Table S2**), strongly suggesting that anaerobic fungi employ GH48 in multi-modular enzymatic complexes to efficiently degrade crystalline cellulose.

This observation is consistent with the powerful degradation activity of fungal multi-modular complexes previously demonstrated by Haitjema et al.,^4^. Although fungal and bacterial dockerins are evolutionary divergent, members of bacterial GH48s have indeed been recognized as the main catalytic component of a processive cellulase in *Clostridium thermocellum* (i.e. *CelS*), exhibiting exo-cellulolytic activity^53^. Albeit at lower protein abundance, peptides also matched fungal cellulosome signature sequences containing carbohydrate binding modules (CBM6 and CBM13) together with GH43s and domains indicative for dockerins (**Table 1**). CBM10, thought to be linked to fungal cellulosomes^21,24,52^, were not detected. Furthermore, in resemblance to the overall metaproteome landscape investigated in this study, the fungal CAZymes had high redundancy across the fungal species as well as high prevalence within each genome.

### Metatranscriptomics of as-yet uncultured populations supports predictions that fungi are active lignocellulose-degraders within the rumen ecosystem

As only a limited number of genomes from anaerobic fungi are currently publicly available, we expected that our RUS-refDB would only represent a fraction of the anaerobic fungal population in the rumen. In an attempt to overcome this constraint, we constructed a complementary database based on a fungal-derived metatranscriptome (‘MT-funDB’), originating from 423,409,432 raw Illumina reads (∼63.5 Gbps) recovered from the fungal community that colonized milled switchgrass during rumen-incubation (**Supplementary Table S3**). After quality filtering and assembly of the raw reads, we identified a total of 4,581,844 expressed genes for which 4,550,231 (99.31%) were predicted to encode proteins. The assembled metatranscriptome was filtered against the genome of the cultured fungi before the MT-funDB was used as a database to identify peptides derived from uncultured rumen fungi within the generated metaproteome data. As with the proteomes of the five fungal isolates, this mapping effort revealed a repertoire consisting of CAZymes belonging to the families of GH3, GH8, GH9, GH10, GH11, GH13 (subfamilies), GH36 and GH48 (**Figure 3, Supplementary Data S2**). While only detected amongst the bacterial population in RUS_refDB, CAZymes were additionally assigned to the families of GH77 and GH94. As fragments believed to be of bacterial origin were observed in MT_funDB, we searched the detected gene sequences of the GH48 representatives against the NR database, which indicated that these protein sequences best resembled glycoside hydrolase family 48 of *Ruminococcus* sp. (sequence similarity ranging from 64 to 97% identity). Despite this seemingly conflicting result, this was not unexpected, given the scarcity of characterized fungal GH48s and the documented frequency of inter-kingdom horizontal gene transfer of catalytic domains in gut ecosystems^4,54^, especially for GH48s^55^. Nevertheless, while we postulate that these active GH48s originate from anaerobic fungi, likely achieved through HGT events, we cannot exclude that bacterial transcripts are present in the metatranscriptome.

### Virome activity in ruminant biomass-degradation

To further enhance our understanding of the role of rumen viruses and how they might shape the different microbial populations within the rumen ecosystem, we analyzed the proteins that were detected in our metaproteomes and that originated from genomic material of viral origin. In accordance with recent research efforts to elucidate the role of the rumen virome, a significant portion of the RUS-refDB proteins (switchgrass: 56; rumen fluid: 62) were assigned to the 913 viral scaffolds we recovered from our switchgrass-associated rumen metagenome^31,56^. Recent studies have demonstrated that some viruses contribute to polysaccharide degradation directly, as they encode glycoside hydrolases^57^, or indirectly through infection of carbohydrate-degrading microorganisms^5^. Accordingly, when mining the genomic content of the viral scaffolds, we identified CAZyme domains within 444 protein-coding genes (**Supplementary Figure S3**). The most prominent was glycoside hydrolase family 25 (58 genes), which contains dominantly enzymes that can hydrolyze the β-1,4-glycosidic bond between N-acetylmuramic acid and N-acetylglucosamine in the carbohydrate backbone of bacterial peptidoglycan and are essential to modify and lyse the bacterial cell wall^58–60^ contributing to intra-ruminal nitrogen turnover. However, none of these viral CAZymes were detected in the metaproteomics data. In general, only a few putative auxiliary metabolic genes were detected within metaproteomes; three bacterial extracellular solute-binding proteins and an oxidoreductase, consistent with a potentially indirect role of viruses in supporting biomass degradation (protein sequences can be found in **Supplementary Text S1)**. Also two ribosomal proteins were found amongst the detected proteins in our data, further reinforcing a recent observation indicating that viruses can modulate the translation upon infection as a strategy to exploit its host^61^. Not surprisingly, a vast majority of the detected viral proteins could not be assigned to any known function, and their purpose in the microbiome cannot be assessed at this time and will require further protein characterization efforts. Several of the detected viral-associated protein groups showed low redundancy and relatively high protein abundance, including a protein detected at the upper range of the protein detection level (average log_2_(LFQ) score = 31.5; gene ID ‘Vir_gene_id_42007’ in **Supplementary Data 1**, the protein sequence can be found in **Supplementary Text S1**). This protein showed high homology (using Phyre^2^: <u>P</u>rotein <u>H</u>omology/Analog<u>Y R</u>ecognition <u>E</u>ngine) to a porter protein, directly involved in the capsid formation and previously found highly abundant in a virion-associated metaproteome^62^. Notably, this protein was detected in the switchgrass fiber fraction samples, yet was absent in the rumen fluid samples. Overall, the numerous viral proteins observed in this study, several quantified at high protein detection level demonstrating their presence and activity, strongly advocate the need for comprehensively studying the rumen virome.

### Towards a holistic understanding of the functional roles of rumen populations

The initial degradation of complex plant fiber makes the carbon pool available for downstream metabolism that encompasses the intricate microbial food web within the rumen, ultimately providing access to otherwise inaccessible nutrients to the host. Concurrent with previous rumen metaproteome and -transcriptome studies^23,63^, our analysis revealed that the prokaryotic population in the rumen plays significant roles in many of the key reactions in the rumen system (**Figure 4**).

**Figure 4:**
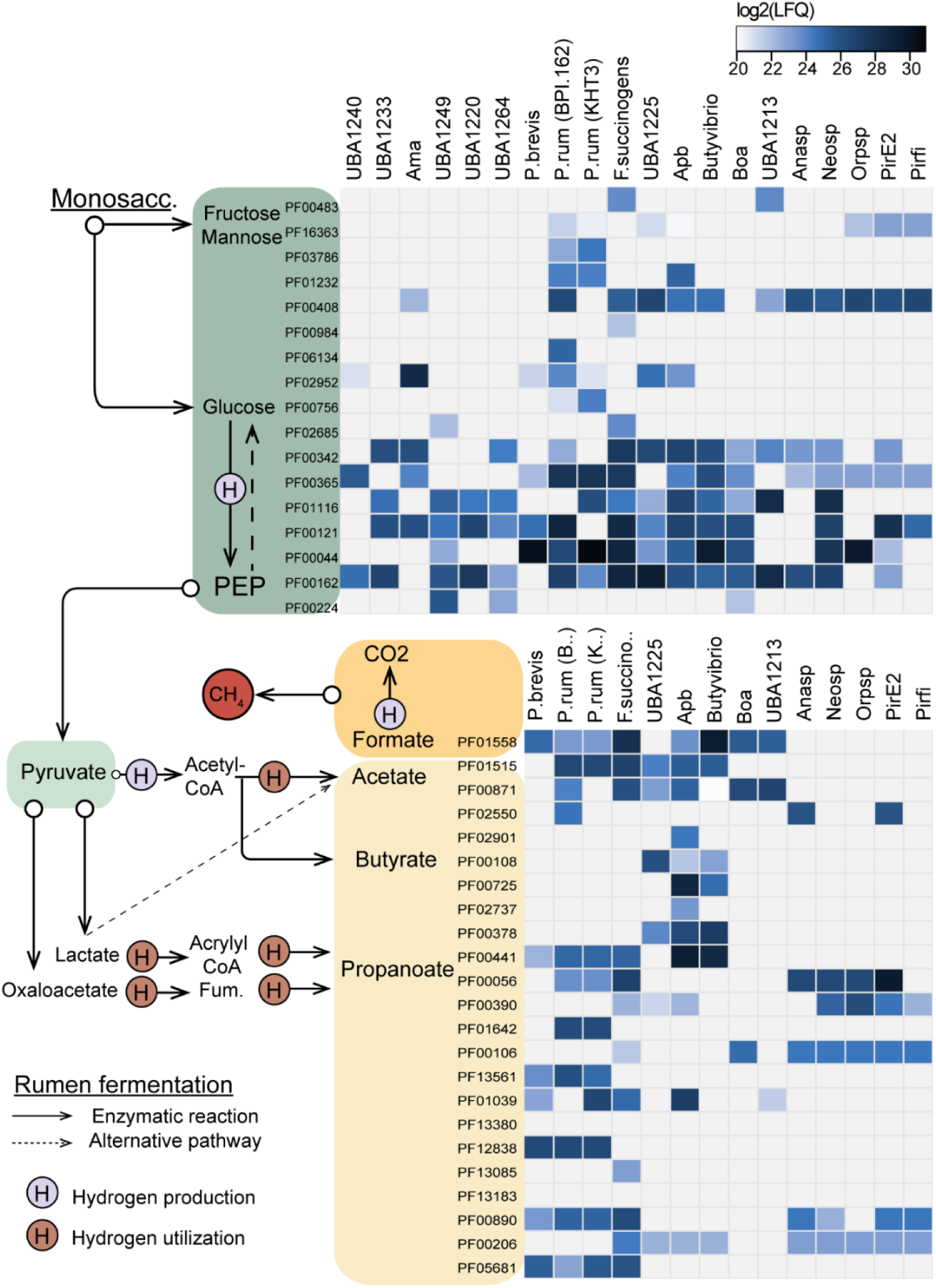
Metabolic reconstruction of key players intermediate rumen fermentation as determined in this study. The heat map shows the detection of proteins associated to main metabolic pathways (listed as pfam IDs) found in the most active genomes/MAGs (indicated on the top: Anasp, *Anaeromyces robustus;* Pirfi, *Piromyces finnis; PirE2, Piromyces* sp. E2; Neosp, *Neocallimastix californiae*; Orpsp; *Orpinomyces* sp.). The colors in the heat map indicates the protein detection levels reported as the average log2(LFQ)-scores for each biological replicate, where light blue represent lower detection levels while darker blue is high protein detection. Only the proteins from the switchgrass are included in the current figure. A comprehensive table including proteins detected in all MAGs/genomes included in RUS-refDB, proteins associated to the rumen fluid and the functional categorization of the pfam IDs can be found in **Supplementary Data S1**.

While glycolysis was, not surprisingly, a widely observed trait across several phylogenetic groups, mannose and fructose metabolism was mainly limited to strains of *P. ruminicola, F. succinogenes* and the uncultured UBA1213. *Prevotella ruminicola* and *F. succinogenes* additionally displayed a relatively protein high detection level of phosphotransacetylase (PF01515) related to acetate production, in addition to several of the key proteins related to the generation of propionate, mainly via oxaloacetate [lactate/malate dehydrogenase (PF00056/PF02866), Methylmalonyl-CoA mutase (PF01642), Acetyl-CoA carboxylase (PF01039)] and fumarate [Succinate dehydogenase/fumarate reductase (PF12838), Fumarase (PF05681)]. As expected due to the close phylogenetic relation to *Butyrivibrio*, genomic content of APb also revealed a metabolic capacity for butyrate production, and its active role in butyrate synthesis in the rumen was supported by the detection of these proteins [Acetyl-CoA acetyltransferase (PF00108), 3-hydroxyacyl-CoA dehydrogenase (PF00725/PF02737), Enoyl-CoA hydratase/isomerase (PF00378), Acyl-CoA dehydrogenase (PF00441)] in the metaproteome data (**Figure 4**).

Although anaerobic fungi have been reported to participate in rumen fermentation, only a few genes related to for example acetate production seem to be “switched on” at the sampling timepoint for our dataset. This may be due to slow growth rates and low protein abundance for these gene sets. Furthermore, while complete glycolysis pathways are annotated for all currently cultivated fungal genomes, only a full set of glycolysis proteins aligning to the genes of *Neocallimastix* was detected at high protein detection levels in our metaproteome data, suggesting that anaerobic fungi only play a minor active role in the downstream carbon flow. Seen in context with the high detection level of fungal enzymes for cellulose decomposition, this emphasizes that a key role of anaerobic fungi at this phase of the biomass degradation (48 hours) is likely to function in recalcitrant fiber degradation of lignin-enriched fiber residues, whereas bacteria encompass a wider functional repertoire, including degradation of more-easily digestible fibers and fermentation.

## Conclusions

While our understanding of the rumen microbiome has increased significantly in recent years, the majority of this knowledge has been restricted to the bacterial population. Insights into the role of anaerobic rumen fungi have been limited to a few studies and still very little is known about the overall ecology of anaerobic rumen fungi as part of the rumen microbiome and their contribution to the biomass-degrading process in the native habitat. In the current study, we report a time dependent scenario within the rumen ecosystem where bacteria appear to have occupied multiple functional niches, while anaerobic fungi seem to dictate the degradation of resilient lignocellulosic plant material. Here, members of the glycoside hydrolase family GH48 were detected at elevated levels and appeared to come nearly exclusively from the rumen fungi. Furthermore, it appears as if the bacterial population in the rumen is primarily involved in degradation of hemicellulose, at least for plant material that has been incubated in the rumen for 48 hours. Overall, these results suggest that anaerobic fungi have a strongly adherent phenotype and colonize recalcitrant plant cell wall material that is likely too large in dimension/particle size to pass out of the rumen. Furthermore, we speculate that their adherent strategy is to maintain their population size in the rumen and prevent them from being washed out, given that they grow slower than the general rumen turnover rate. Although these results broaden our understanding of the native function of anaerobic rumen fungi, spatial and temporal experiments would certainly be beneficial to provide further support of the hypothesis that the detected proteins are ubiquitously involved in the degradation of recalcitrant biomass in the rumen and are essential to the nutrition and well-being of their host animal.

## Material and Methods

### Rumen incubation and sample collection

Air-dried switchgrass was milled to pass through a 2 mm sieve and weighed into individual *in situ* nylon bags (50 μm pores; Ankom Technology, Macedon, NY, USA). To enrich for lignocellulolytic microorganisms, the Nylon bags, each containing 5 g of air-dried switchgrass, were placed in the rumen of two cannulated cows as described previously (Hess et al. 2011). Nylon bags were retrieved from the cow’s rumen after 48 h, washed immediately with PBS buffer (pH7) to remove loosely adherent microbes, frozen immediately in liquid nitrogen and transported to the laboratory. Samples were stored at -80°C until protein and RNA extraction was performed. All animal procedures were performed in accordance with the Institution of Animal Care and Use Committee (IACUC) at the University of Illinois, under protocol number #06081.

### Construction of a rumen-specific reference database (RUS-refDB)

A collection of protein sequences from rumen associated microorganisms was generated from a total of 122 microbial genomes (from MAGs and isolates) and 931 metagenome-assembled viral scaffolds. To account for the prokaryotic rumen population and their major metabolic function we selected 12 genomes from the Hungate1000 project^6^. We supplemented these core genomes with the genome of *Fibrobacter succinogenes* S85^32^ and *Methanobrevibacter ruminantium* M1^33^. To reduce cultivation bias, the sequence database was composed of metagenome assembled genomes (MAGs), originating from Hess et al. (2011) as well as a recent re-assembly of this metagenome published by Parks et al. (2017). Genome redundancy was reduced by removing genomes with an amino-acid identity (AAI) > 99% (CompareM v.0.0.13), of which the MAGs with the highest quality (CheckM v.1.0.18) were kept for downstream analysis. This resulted in a non-redundant catalogue of high-quality MAGs, composed of 7 and 96 MAGs from Hess et al. (2011) and Donovan et al. (2017), respectively. Genes in metagenome-assembled viral scaffolds, previously recovered from a rumen metagenome^30,31^, were predicted with GeneMark^64^. In order to elucidate the functional roles of anaerobic fungi in the rumen, we also included protein sequences from the genomes of the five cultivated anaerobic fungi available at the time; *Anaeromyces robustus, Neocallimastix californiae* G1, *Orpinomyces* sp., *Piromyces* sp. E2 and *Piromyces finnis* downloaded from MycoCosm^65^ (available from https://mycocosm.jgi.doe.gov). A summary of the MAGs, MAVSs, and SAGs that made up our reference database is provided in **Table S1**. This sequence collection was further used as a comprehensive reference database (“RUS-refDB”) for mapping of the metaproteome data, as described below.

### Phylogenetic tree

For the phylogenetic tree we searched each genome and MAG included in the RUS-refDB for 21 ribosomal proteins (L1, L3, L4, L5, L6, L11, L13, L18, L22, L24, S2, S5, S8, S9, S10, S11, S12, S13, S15, S17 and S19). The resulting ribosomal protein sequences were aligned separately using MUSCLE^66^ v3.8.31 and manually checked for duplication and misaligned sequences. For further alignment clean-up, GBlocks^67^ v.0.91b with a relaxed selection of blocks (Gblocs settings: -b2=50 -b3=20 -b4=2 -b5=a) was employed. The alignments were then concatenated using catfasta2phyml.pl (https://github.com/nylander/catfasta2phyml) with the parameter ‘-c’ to replace missing ribosomal proteins with gaps (-). The initial maximum likelihood phylogenetic tree was constructed using RAxML^68^ v.8.2.12 (raxmlHPC-SSE3 under PROTGAMMA with WAG substitution matrix and 100 rapid bootstrap inferences). One MAG (UBA1267) was not included in the ribosomal protein tree due to undetermined values. A complete version of this tree is available in Newick format as **Supplementary Data S3**. This tree was then re-built from a separate alignment including two ribosomal proteins (L3 and S9) from the five rumen fungi included in RUS-refDB, and finally visualized using iTol^69^.

### Metaproteomics – protein extraction and mass spectrometry

Protein extraction and mass spectrometry were performed on rumen-incubated switchgrass as described previously in Naas et al. (2018). In brief, proteins were extracted from bulk rumen fluid and different fractions of the solid rumen-incubated biomass. Solid biomass was ground using a Biopulverizer (Biospec, Bartlesville, OK) and liquid nitrogen. SIGMAFAST protease inhibitor was added to prevent protein degradation during sample preparation. Protein concentrations were determined using the bicinchoninic acid (BCA) protein assay (ThermoFisher Pierce, Waltham, MA). Urea and dithiothreitol (DTT) were added to all samples to a final concentration of 8 M and 10 mM, respectively and incubated at 60°C for 30 minutes to denature and reduce proteins. Protein digestion was performed at 37°C (235 rpm) for 3 hours after CaCl_2_ trypsin was added to a 1 mM final concentration and in a 1:50 trypsin:protein (w/w) ratio, respectively. After sample clean-up and concentration, samples were analyzed by reversed phase LC-MS/MS using a Waters nanoACQUITY™ UPLC system (Millford, MA) coupled with an Orbitrap Velos mass spectrometer (Thermo Fisher Scientific, San Jose, CA). The obtained MS/MS scans were subsequently analyzed using MaxQuant^70^ v.1.6.0.13, and proteins quantified using the MaxLFQ^71^ algorithm implemented in MaxQuant. Peptides were identified by searching the MS/MS datasets against the reference databases. To identify common contaminants introduced during sample preparation, this database was complemented with common contaminants, such as human keratin and bovine serum albumin, as well as with reversed sequences in order to estimate the false discovery rate. Tolerance levels for peptide identifications were 6 ppm and 0.5 Da for MS and MS/MS, respectively, and two missed cleavages of trypsin were allowed. Carbamidomethylation of cysteine residues was used as a fixed modification, while oxidation of methionines and protein N-terminal acetylation were used as variable modifications. All identifications were filtered in order to achieve a protein false discovery rate of 1% using the target-decoy strategy. The software Perseus version 1.6.0.7^72^ was used for downstream interpretation and quality filtering, including removal of decoy database hits, hits only identified by site and contaminants. Finally, at least one unique peptide per protein was required for a protein to be considered as valid.

### Metatranscriptomics - total RNA extraction and Poly(A) mRNA purification

Total RNA was isolated as described previously^73^. First, frozen rumen-incubated biomass (switchgrass) was manually ground to powder in the presence of liquid nitrogen and immediately added to TRIzol reagent (Invitrogen, Carlsbad, CA). Next, the biomass/TRIzol mixture was transferred into a 2 mL microcentrifuge tube containing Lysing Matrix E (MP Biomedicals Solon, OH), followed by bead beating (3 x 1 min at room temperature, 2 min at 4°C between individual beating steps) using a Mini-Beadbeater-16 (Biospec Products, Bartlesville OK). Homogenized samples were centrifuged (12,000 x *g*, 10 min at 4°C); the supernatant was transferred to new tubes and incubated at room temperature for 5 min. Subsequent TRIzol-based RNA isolation was performed according to manufacturer’s instructions. Poly(A) mRNA was isolated from total RNA with MicroPoly(A)Purist kit (Invitrogen, Carlsbad, CA) following the manufacturer’s instructions.

The prepared libraries were quantified using KAPA Biosystem’s next-generation sequencing library qPCR kit and run on a Roche LightCycler 480 real-time PCR instrument. The quantified libraries were then multiplexed, and the library pool was then prepared for sequencing on the Illumina HiSeq platform utilizing a TruSeq paired-end cluster kit, v3, and Illumina’s cBot instrument to generate a clustered flow cell. Sequencing was performed on the Illumina HiSeq2000 using a TruSeq SBS sequencing kit, v3, following a 2−150 indexed run recipe. Adapter sequences and low-quality reads (Q < 10) were trimmed and the reads were further filtered to remove process artifacts using BBDuk included in BBTools^74^ from JGI. After trimming and filtering, human and ribosomal RNA reads were removed by mapping sequences against a modified Silva database^75^ using BBMap^74^. Cleaned reads were combined and the metatransciptome was assembled using MEGAHIT^76^ v.0.2.0. The transcripts were then mapped against the assembled genome of each of the five fungal species represented in RUS-refDB (i.e. *Anaeromyces robustus, Neocallimastix californiae* G1, *Orpinomyces* sp., *Piromyces* sp. E2 and *Piromyces finnis)* using BWA-MEM^77^, and those aligned to genomes were excluded from downstream analysis. Additionally, contigs shorter than 1 kb were removed from the dataset. TransDecoder v.2.0.1 with default settings was used to identify open reading frames (ORFs) within the transcripts and the resulting sequences (256 232 ORFs) was used as a fungal-associated database (“MT-funDB”). The MS scans retrieved from the extracted metaproteome was then searched against fun-DB in the same manner as described previously for RUS-refDB. A comprehensive table of detected proteins in switchgrass fiber and rumen fluid for each genome/MAGs can be found in **Supplementary Data S2.**

### Functional annotation and metabolic reconstruction

All protein sequences included in RUS-refDB and MT-funDB were functionally annotated using InterProScan5^78^ v.5.25-64, including search against pfam and CDD databases, Gene Ontology (GO) annotation and mapping to KEGG pathway information. CAZymes in RUS-refDB were additionally annotated using the CAZy annotation pipeline^79^. This functional annotation information was added to the detected protein groups in Perseus, and manually searched for specific metabolism. To ensure high confidence results in the reported CAZyme and cellulosomal signature sequences, the protein had to be detected in both biological replicates (i.e. in both cows) in at least one of the two samples (i.e. switchgrass fiber and rumen fluid). Protein groups not fulfilling these criteria were omitted from the main results. For the reconstruction of active pathways involving monosaccharide degradation and fermentation, we scanned the detected protein in each of the annotated genomes and MAGs for signature pfam IDs, and further validated its function using its Interpro, CDD and GO annotation. A complete or nearly complete set of pathway genes needed to be turned on for a genome to be considered as actively involved in a respective metabolism. The protein detection levels of each protein group are reported as the average Log_2_(LFQ) for each biological replicate, which enables the quantification of the active metabolic function of the keystone rumen populations. Heat maps were generated with the ggplots package heatmap.2 in RStudio v.3.6.1^80^. Furthermore, only the protein profile for switchgrass fiber is displayed in the constructed CAZyme and metabolic heat maps in order to reduce complexity. A comprehensive table of detected proteins in switchgrass fiber and rumen fluid for each genome/MAG can be found in **Supplementary Data S1.**

## Data availability

The mass spectrometry proteomics data have been deposited to the ProteomeXchange Consortium (http://proteomecentral.proteomexchange.org) via the PRIDE^81^ partner repository with the dataset identifier PXD017007. The metatranscriptome raw files are submitted to NCBI SRA, accession numbers SRR9001933, SRR6230176, SRR6230410, SRR9001942, SRR9002087 and SRR6230409. The references for the genomes, metagenome-assembled genomes and viral scaffolds are listed in **Supplementary Table S1**.

## Acknowledgements

A portion of the research was performed using EMSL, a DOE Office of Science User Facility sponsored by the Office of Biological and Environmental Research and located at Pacific Northwest National Laboratory. The work conducted by the U.S. Department of Energy Joint Genome Institute, a DOE Office of Science User Facility, is supported by the Office of Science of the U.S. Department of Energy under Contract No. DE-AC02-05CH11231. We are grateful for support from The Research Council of Norway (FRIPRO program, P.B.P. and L.H.H.: 250479), as well as the European Research Commission Starting Grant Fellowship (awarded to P.B.P.; 336355 - MicroDE).

## Authors contributions

M.H. conceived and designed the experiments. H.P., R.I.M. and M.H. performed the experiments. L.H.H., C.B.G., C.S., A.N.D., H.P., M.Ø.A., H.B., A.C., N.I., S.T., L.P., P.B.P. and M.H. generated and analyzed the data. L.H.H., P.B.P. and M.H. wrote the major part of the manuscript. L.H.H., C.G.B., C.S., A.D.N., H.P., M.Ø.A., H.B., A.C., S.R., V.L., B.H., M.A.O., I.V.G., S.T., R.I.M., L.P., P.B.P. and M.H. contributed to the final version of the manuscript.

