## Supplementary Material for "Proteome specialization of anaerobic fungi during ruminal degradation of recalcitrant plant fiber"

*List of content:*

**Figure S1**: Scatterplots of the proteins quantified from the switchgrass fiber (SF) and rumen fluid (RF). The Pearson coefficient is indicated for each scatterplot.

**Figure S2:** Quantitative breakdown of detected protein groups annotated as CAZyme when searching the metaproteome against the 'RUS-refDB' database.

**Figure S3:** Predicted CAZymes in the genetic content on the rumen virome.

**Table S1**: Overview of the genomes, metagenome-assembled genomes and the scaffolds used to generate RUS-refDB.

**Table S2**: Extended table of detected protein groups associated with cellulosome signature domains (dockerins).

**Table S3:** Metatranscriptome Quality Filtering Stats.

**Text S1:** FASTA sequences of the putative auxiliary proteins, detected within the metaproteome. This list also includes one additional viral protein sequence detected at the upper range of the protein detection level.

*In separate files:*

**Supplementary Data S1:** Extended table of detected protein groups mapping against RUS-refDB

**Supplementary Data S2:** Extended table of detected protein groups annotated as CAZymes mapping against MT-funDB.

**Supplementary Data S3**: The complete concatenated ribosomal protein tree in Newick format.

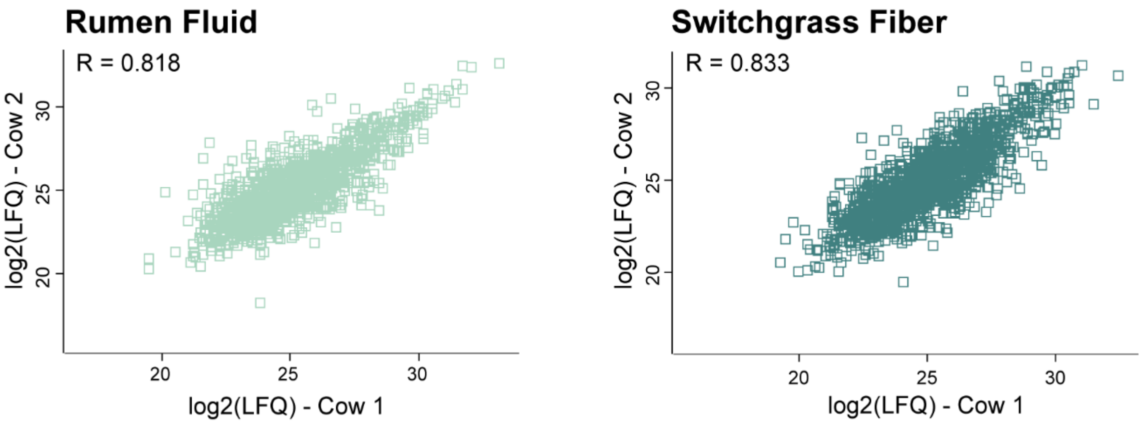


**Figure S1**: Correlation analysis of $\log_2$(LFQ) values between the two biological replicates (Cow 1 and Cow 2) in the rumen fluid and switchgrass fiber microhabitats. The Pearson coefficient for each comparison is shown in each plot.

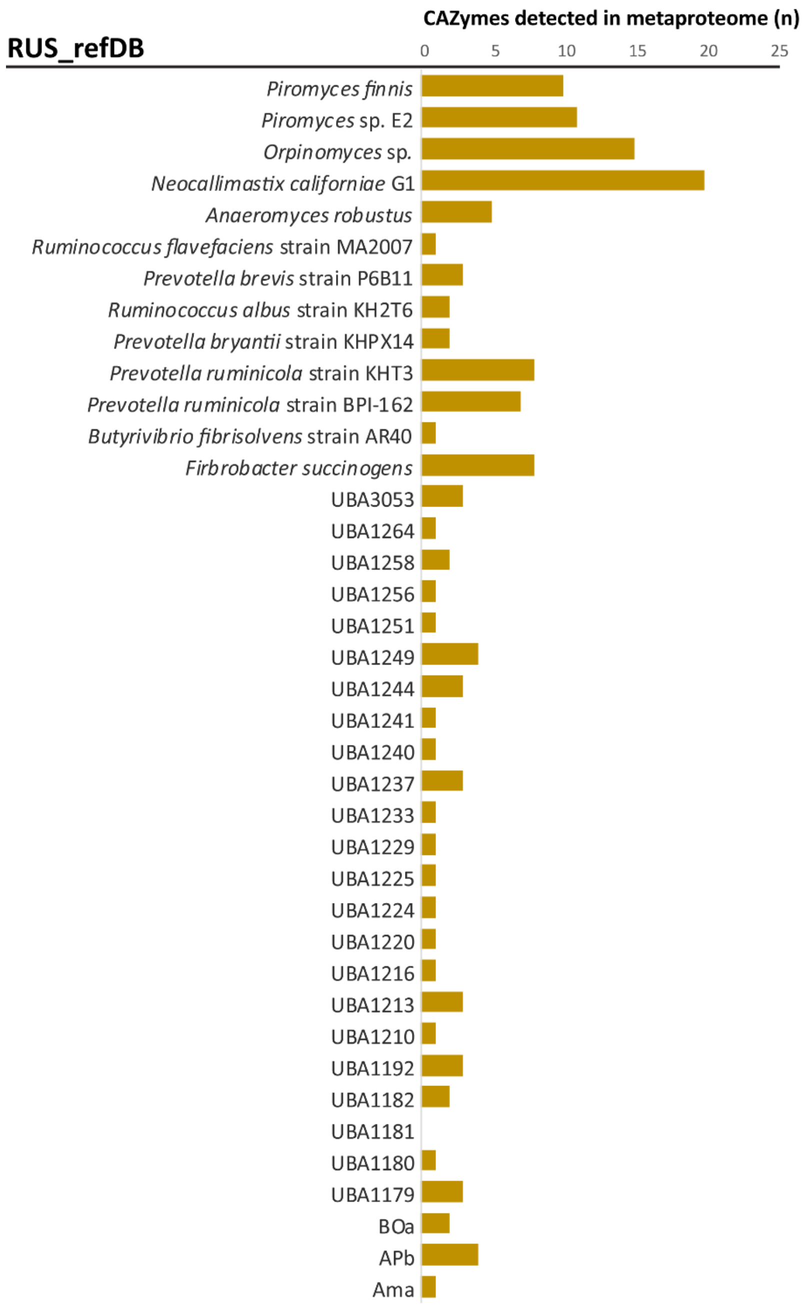


**Figure S2:** Quantitative breakdown of detected protein groups annotated as CAZyme (includes GHs, CEs, GTs, CBMs and dockerins) when searching the metaproteome against the 'RUS-refDB' database. Only proteins detected in both animals in at least one of the two microhabitats were included. Moreover, if a protein group consisted of more than one protein identification, only the first identification was considered.

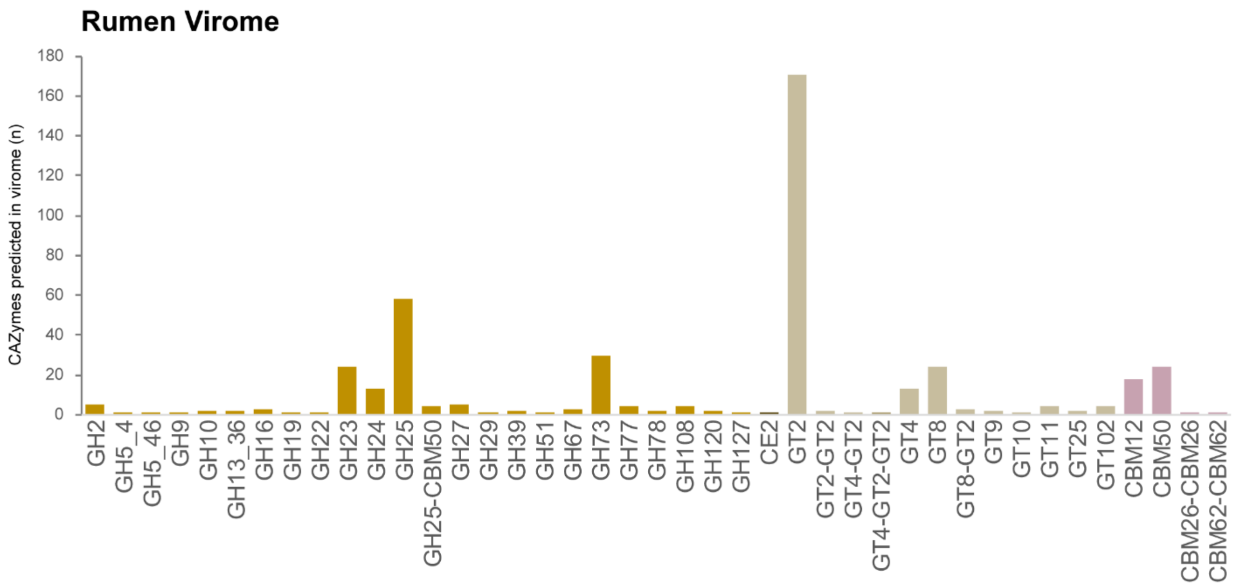


**Figure S3:** Predicted CAZymes in the genetic content of the rumen virome, colored by enzyme classes (GH, Glycoside Hydrolases; CE, Carbohydrate Esterases, GT, Glycosyl Transferases; CBM; Carbohydrate-Binding Modules). None of these were detected in the metaproteomics data.

**Table S1**: Overview of the genomes, metagenome-assembled genomes and the virome used to generate RUS-refDB. This table includes the number of predicted genes, as well as the number of detected proteins in the rumen fluid (RF) and switchgsrass fraction (SF) when mapped against the metaproteomics data.

| **IDs in MP** | **Organism Name** | **Ref.** | **Predicted Genes (n)** | **Detected proteins (n)** | |
|---|---|---|---|---|---|
| | | | | ***RF*** | ***SF*** |
| ***Genomes of anaerobic cultivated fungi*** | | | | | |
| *jgi\|Anasp1* | Anaeromyces robustus v1.0 | [4] | 12832 | 171 | 341 |
| *jgi\|Neosp1* | Neocallimastix californiae G1 v1.0 | [4] | 20219 | 360 | 787 |
| *jgi\|Orpsp1_1* | Orpinomyces sp. | [18] | 18936 | 121 | 320 |
| *jgi\|PirE2_1* | Piromyces sp. E2 v1.0 | [4] | 14648 | 172 | 329 |
| *jgi\|Pirfi3* | Piromyces finnis v3.0 | [4] | 10992 | 172 | 316 |
| ***MAGs from a rumen metagenome*** | | | | | |
| *AH* | AH [Bacteroidales] | [31] | 3652 | 53 | 60 |
| *AMa* | AMa [Spirochaetales] | [31] | 3103 | 47 | 82 |
| *AN* | AN [Clostridiales] | [31] | 3967 | 29 | 55 |
| *APb* | APb [Clostridiales] | [31] | 4239 | 97 | 237 |
| *AQ* | AQ [Bacteroidales] | [31] | 3151 | 85 | 50 |
| *AS1a* | AS1a [Clostridiales] | [31] | 2302 | 15 | 11 |
| *BOa* | BOa [Clostridiales] | [31] | 4188 | 121 | 126 |
| *UBA3053* | UBA3053 [Bacteroidales bacterium] | [30] | 1694 | 39 | 32 |
| *UBA3054* | UBA3054 [Candidatus UBP3 bacterium] | [30] | 1217 | 4 | 3 |
| *UBA1173* | UBA1173 [Bacteroidales bacterium] | [30] | 1992 | 48 | 54 |
| *UBA1174* | UBA1174 [Elusimicrobia bacterium] | [30] | 1248 | 4 | 5 |
| *UBA1175* | UBA1175 [Lachnoclostridium sp.] | [30] | 2287 | 35 | 51 |
| *UBA1176* | UBA1176 [Bacteroidales bacterium] | [30] | 2144 | 37 | 38 |
| *UBA1177* | UBA1177 [Candidatus UBP6 bacterium] | [30] | 1931 | 3 | 5 |
| *UBA1178* | UBA1178 [Bacteroidales bacterium] | [30] | 1410 | 19 | 20 |
| *UBA1179* | UBA1179 [Bacteroidales bacterium] | [30] | 1699 | 31 | 35 |
| *UBA1180* | UBA1180 [Bacteroidales bacterium] | [30] | 1629 | 43 | 31 |
| *UBA1181* | UBA1181 [Bacteroidales bacterium] | [30] | 2604 | 55 | 85 |
| *UBA1182* | UBA1182 [Bacteroidales bacterium] | [30] | 2090 | 28 | 31 |
| *UBA1183* | UBA1183 [Verrucomicrobia bacterium] | [30] | 1000 | 1 | 0 |
| *UBA1184* | UBA1184 [Bacteroidales bacterium] | [30] | 2301 | 57 | 78 |
| *UBA1185* | UBA1185 [Alphaproteobacteria bacterium] | [30] | 792 | 3 | 0 |
| *UBA1186* | UBA1186 [Cyanobacteria bacterium] | [30] | 1625 | 17 | 23 |
| *UBA1187* | UBA1187 [Bacteroidales bacterium] | [30] | 1477 | 61 | 35 |
| *UBA1188* | UBA1188 [Bacillales bacterium] | [30] | 1004 | 7 | 6 |
| *UBA1189* | UBA1189 [Bacteroidales bacterium] | [30] | 2498 | 36 | 41 |
| *UBA1190* | UBA1190 [Ruminococcus flavefaciens sp.] | [30] | 2946 | 40 | 50 |
| *UBA1191* | UBA1191 [Firmicutes bacterium] | [30] | 1316 | 23 | 35 |
| *UBA1192* | UBA1192 [Bacteroidales bacterium] | [30] | 1833 | 32 | 29 |
| *UBA1193* | UBA1193 [Mageeibacillus sp.] | [30] | 2111 | 18 | 25 |
| *UBA1194* | UBA1194 [Clostridiales bacterium] | [30] | 1990 | 12 | 21 |
| *UBA1195* | UBA1195 [Bacteroidales bacterium] | [30] | 1997 | 20 | 22 |

| | | | | | |
|---|---|---|---|---|---|
| *UBA1196* | UBA1196 [Candidatus UBP6 bacterium] | [30] | 1693 | 3 | 4 |
| *UBA1197* | UBA1197 [Bacteroidales bacterium] | [30] | 1802 | 18 | 16 |
| *UBA1198* | UBA1198 [Faecalibacterium sp.] | [30] | 1299 | 8 | 10 |
| *UBA1199* | UBA1199 [Bacteroidales bacterium] | [30] | 1191 | 21 | 22 |
| *UBA1200* | UBA1200 [Verrucomicrobia bacterium] | [30] | 2626 | 9 | 11 |
| *UBA1201* | UBA1201 [Lachnospiraceae bacterium] | [30] | 2105 | 27 | 37 |
| *UBA1202* | UBA1202 [Elusimicrobia bacterium] | [30] | 1288 | 5 | 6 |
| *UBA1203* | UBA1203 [Myxococcales bacterium] | [30] | 2161 | 0 | 12 |
| *UBA1204* | UBA1204 [Bacteroidales bacterium] | [30] | 2202 | 32 | 31 |
| *UBA1205* | UBA1205 [Bacteroidales bacterium] | [30] | 2054 | 26 | 26 |
| *UBA1206* | UBA1206 [Clostridiales bacterium] | [30] | 1747 | 17 | 27 |
| *UBA1207* | UBA1207 [Ruminococcaceae bacterium] | [30] | 1622 | 13 | 14 |
| *UBA1208* | UBA1208 [Bacteroidales bacterium] | [30] | 2405 | 61 | 36 |
| *UBA1209* | UBA1209 [Candidatus UBP6 bacterium] | [30] | 1647 | 5 | 8 |
| *UBA1210* | UBA1210 [Ruminococcus flavefaciens sp.] | [30] | 2272 | 44 | 68 |
| *UBA1211* | UBA1211 [Verrucomicrobia bacterium] | [30] | 2824 | 10 | 12 |
| *UBA1212* | UBA1212 [Mageeibacillus sp.] | [30] | 2189 | 22 | 23 |
| *UBA1213* | UBA1213 [Ruminococcaceae bacterium] | [30] | 2202 | 137 | 113 |
| *UBA1214* | UBA1214 [Catabacter sp.] | [30] | 1758 | 17 | 17 |
| *UBA1215* | UBA1215 [Bacteroidales bacterium] | [30] | 1559 | 22 | 22 |
| *UBA1216* | UBA1216 [Bacteroidales bacterium] | [30] | 1810 | 36 | 23 |
| *UBA1217* | UBA1217 [Bacteroidales bacterium] | [30] | 1792 | 47 | 28 |
| *UBA1218* | UBA1218 [Alphaproteobacteria bacterium] | [30] | 812 | 1 | 3 |
| *UBA1219* | UBA1219 [Bacteroidales bacterium] | [30] | 2281 | 43 | 36 |
| *UBA1220* | UBA1220 [Succinivibrionaceae bacterium] | [30] | 1140 | 43 | 61 |
| *UBA1221* | UBA1221 [Cyanobacteria bacterium] | [30] | 2139 | 11 | 11 |
| *UBA1222* | UBA1222 [Dehalococcoidia bacterium] | [30] | 815 | 5 | 5 |
| *UBA1223* | UBA1223 [Bacteroidales bacterium] | [30] | 1650 | 39 | 32 |
| *UBA1224* | UBA1224 [Catabacter sp.] | [30] | 2733 | 28 | 50 |
| *UBA1225* | UBA1225 [Pseudobutyrivibrio sp.] | [30] | 2646 | 52 | 103 |
| *UBA1227* | UBA1227 [Ruminococcaceae bacterium] | [30] | 977 | 11 | 22 |
| *UBA1228* | UBA1228 [bacterium] | [30] | 1943 | 18 | 21 |
| *UBA1229* | UBA1229 [Bacteroidales bacterium] | [30] | 2972 | 37 | 66 |
| *UBA1230* | UBA1230 [Oscillospiraceae bacterium] | [30] | 1402 | 18 | 27 |
| *UBA1231* | UBA1231 [Bacillales bacterium] | [30] | 1279 | 7 | 10 |
| *UBA1232* | UBA1232 [Bacteroidales bacterium] | [30] | 1771 | 16 | 15 |
| *UBA1233* | UBA1233 [Treponema sp.] | [30] | 2880 | 37 | 62 |
| *UBA1234* | UBA1234 [Clostridiales bacterium] | [30] | 1456 | 4 | 9 |
| *UBA1235* | UBA1235 [Ca. Saccharibacteria bacterium] | [30] | 603 | 0 | 0 |
| *UBA1236* | UBA1236 [Ruminococcaceae bacterium] | [30] | 1514 | 14 | 28 |
| *UBA1237* | UBA1237 [Bacteroidales bacterium] | [30] | 2153 | 22 | 25 |
| *UBA1238* | UBA1238 [Myxococcales bacterium] | [30] | 3136 | 8 | 15 |
| *UBA1239* | UBA1239 [Clostridiales bacterium] | [30] | 1354 | 5 | 10 |
| *UBA1240* | UBA1240 [Treponema sp.] | [30] | 1972 | 15 | 41 |

| | | | | | |
|---|---|---|---|---|---|
| *UBA1241* | UBA1241 [Lachnoclostridium sp.] | [30] | 2012 | 26 | 39 |
| *UBA1242* | UBA1242 [Catabacter sp.] | [30] | 1070 | 3 | 8 |
| *UBA1243* | UBA1243 [Mageeibacillus sp.] | [30] | 1518 | 14 | 24 |
| *UBA1244* | UBA1244 [Ruminococcaceae bacterium] | [30] | 2075 | 71 | 65 |
| *UBA1245* | UBA1245 [Bacillales bacterium] | [30] | 628 | 0 | 4 |
| *UBA1246* | UBA1246 [Clostridiales bacterium] | [30] | 1884 | 11 | 12 |
| *UBA1247* | UBA1247 [Candidatus UBP3 bacterium] | [30] | 1250 | 2 | 3 |
| *UBA1248* | UBA1248 [Clostridiales bacterium] | [30] | 1983 | 9 | 14 |
| *UBA1249* | UBA1249 [Gammaproteobacteria bacterium] | [30] | 1377 | 127 | 74 |
| *UBA1250* | UBA1250 [Porphyromonadaceae bacterium] | [30] | 1614 | 16 | 15 |
| *UBA1251* | UBA1251 [Lachnospiraceae bacterium] | [30] | 1785 | 33 | 55 |
| *UBA1252* | UBA1252 [Catabacter sp.] | [30] | 1212 | 6 | 13 |
| *UBA1253* | UBA1253 [Bacteroidales bacterium] | [30] | 1677 | 26 | 16 |
| *UBA1254* | UBA1254 [Alphaproteobacteria bacterium] | [30] | 851 | 4 | 0 |
| *UBA1255* | UBA1255 [Mageeibacillus sp.] | [30] | 2126 | 12 | 16 |
| *UBA1256* | UBA1256 [Bacteroidales bacterium] | [30] | 1352 | 39 | 21 |
| *UBA1257* | UBA1257 [Mageeibacillus sp.] | [30] | 1446 | 14 | 18 |
| *UBA1258* | UBA1258 [Lachnospiraceae bacterium] | [30] | 2042 | 26 | 48 |
| *UBA1259* | UBA1259 [Catabacter sp.] | [30] | 1179 | 9 | 12 |
| *UBA1260* | UBA1260 [Catabacter sp.] | [30] | 1179 | 12 | 11 |
| *UBA1261* | UBA1261 [Bacillales bacterium] | [30] | 1194 | 4 | 5 |
| *UBA1262* | UBA1262 [Catabacter sp.] | [30] | 716 | 12 | 8 |
| *UBA1263* | UBA1263 [Verrucomicrobia bacterium] | [30] | 1911 | 9 | 8 |
| *UBA1264* | UBA1264 [Succinatimonas sp.] | [30] | 1152 | 54 | 44 |
| *UBA1265* | UBA1265 [Spirochaetales bacterium] | [30] | 1377 | 9 | 12 |
| *UBA1266* | UBA1266 [Bacteroidales bacterium] | [30] | 1557 | 10 | 8 |
| *UBA1267* | UBA1267 [Lachnospiraceae bacterium] | [30] | 1424 | 14 | 16 |
| ***Reference bacterial/archaeal genomes*** | | | | | |
| *Ga0070636* | Prevotella ruminicola strain BPI-162 | [6] | 3012 | 213 | 178 |
| *Ga0104360* | P. ruminicola strain KHT3 | [6] | 3446 | 205 | 173 |
| *T496DRAFT* | P. brevis strain P6B11 | [6] | 2510 | 168 | 129 |
| *Ga0104364* | P. bryantii strain KHPX14 | [6] | 2707 | 77 | 64 |
| *Ga0104370* | Ruminococcus albus strain KH2T6 | [6] | 3513 | 62 | 81 |
| *Ga0066885* | R. flavefaciens strain Y1 | [6] | 3199 | 70 | 90 |
| *T497DRAFT* | R. flavefaciens strain MA2007 | [6] | 2907 | 75 | 106 |
| *T488DRAFT* | R. flavefaciens strain ND2009 | [6] | 3137 | 63 | 78 |
| *IE37DRAFT* | R. flavefaciens strain SAb67 | [6] | 3384 | 67 | 79 |
| *Ga0070258* | Bacillus licheniformis strain VTM3R78 | [6] | 4158 | 12 | 9 |
| *Ga0070254* | Butyrivibrio fibrisolvens strain AR40 | [6] | 4211 | 46 | 96 |
| *WP_012819846.1* | Fibrobacter succinogens subsp. Succ. S85 | [32] | 3188 | 210 | 349 |
| *WP_012954809.1* | Methanobrevibacter ruminantium strain M1 | [33] | 2135 | 39 | 51 |
| *Ga0117888_1000011* | Rumen methanol enrichment MEC1 | [6] | 44933 | 38 | 44 |
| ***Viral content from a rumen metagenome*** | | | | | |
| *Vir* | ViralScaffolds-RumenMG | [31,55] | 74440 | 62 | 56 |

**Table S2**: Extended table of detected protein groups associated with cellulosome signature domains (dockerins). Protein detection level is given as $log_2$(LFQ) for each biological replicate in samples from rumen fluid (RF) and switchgrass fiber (SF). The numbers in the left marks the proteins (in bold) included in **Table 1**. Only the first protein identification per protein group is shown in this table. Protein groups with more than one protein sequence identification are indicated with *. A complete list can be found in **Supplementary Data S1**.

| | **MAG/genome**<br>**[majority protein ID]** | **CAZyme module** | **Protein detection [$log_2$(LFQ)]** | | | |
|---|---|---|---|---|---|---|
| | | *Microhabitat:*<br>*Cow:* | **RF**<br>**1** | **RF**<br>**2** | **SF**<br>**1** | **SF**<br>**2** |
| *1* | R. flavefaciens strain MA2007<br>[T497DRAFT_00845]* | **CBM4-GH9-DOC1** | 25.1 | 24.6 | 24.1 | 25.0 |
| | R. flavefaciens strain MA2007<br>[T497DRAFT_00649] | DOC1 | 23.3 | NaN | NaN | NaN |
| | R. flavefaciens strain MA2007<br>[T497DRAFT_01962]* | DOC1 | NaN | NaN | NaN | 20.6 |
| | R. flavefaciens strain MA2007<br>[T497DRAFT_01936]* | GH11<br>GH11-CBM22-DOC1-CE1<br>GH11-CE1-DOC1-CE4<br>GH11-GH10<br>GH11-GH11-GH10-DOC1-GH11-CE4 | NaN | NaN | NaN | 22.8 |
| | R. flavefaciens strain MA2007<br>[T497DRAFT_01436] | GH11-CBM22-GH10-DOC1-GH11-CE4 | NaN | 22.0 | NaN | NaN |
| | R. flavefaciens strain MA2007<br>[T497DRAFT_00844] | GH9-CBM3-DOC1 | NaN | NaN | NaN | 21.3 |
| | R. flavefaciens strain Y1<br>[Ga0066885_11723] | DOC1 | NaN | 23.8 | NaN | 23.3 |
| | R. flavefaciens strain Y1<br>[Ga0066885_10689]* | GH11-CBM22-GH10-DOC1-CBM22-CE1<br>GH11-CBM22-GH10-DOC1-GH11-CE4 | NaN | NaN | NaN | 22.5 |
| | UBA1190<br>[UBA1190_contig_427_14] | DOC1 | 22.9 | NaN | NaN | NaN |
| | UBA1190<br>[UBA1190_contig_96797_16] | DOC1 | NaN | NaN | NaN | 19.4 |
| | UBA1210<br>[UBA1210_contig_2415_7] | DOC1 | NaN | NaN | 21.4 | NaN |
| | UBA1210<br>[UBA1210_contig_2415_8] | DOC1 | NaN | NaN | 24.7 | 23.9 |
| | UBA1210<br>[UBA1210_contig_56792_10]* | GH11-CBM22-GH10-DOC1<br>GH11-CBM22-GH10-DOC1-CE1 | NaN | NaN | NaN | 22.7 |
| | Anaeromyces robustus v1.0<br>[jgi\|Anasp1\|296357]* | CBM35-GH26-DOC2-DOC2<br>CBM35-GH26-DOC2-DOC2-DOC2-DOC2-DOC2 | NaN | NaN | NaN | 20.8 |
| | Anaeromyces robustus v1.0<br>[jgi\|Anasp1\|327659]* | CE15<br>CE15-DOC2-DOC2<br>GH11-CBM6-CE15 | NaN | NaN | NaN | 24.4 |
| | Anaeromyces robustus v1.0<br>[jgi\|Anasp1\|294808]* | GH3-DOC2-DOC2-DOC2 | NaN | NaN | 23.5 | NaN |
| *2* | Anaeromyces robustus v1.0<br>[jgi\|Anasp1\|287068]* | **GH43**<br>**GH43-CBM6**<br>**GH43-CBM6-CBM13-DOC2-DOC2**<br>**GH43-CBM6-DOC2-DOC2** | NaN | NaN | 23.9 | 24.0 |
| | Anaeromyces robustus v1.0<br>[jgi\|Anasp1\|327938] | GH43-CBM6-DOC2-DOC2 | NaN | 20.8 | NaN | NaN |
| | Anaeromyces robustus v1.0<br>[jgi\|Anasp1\|233234]* | GH48-DOC2-DOC2 | NaN | NaN | NaN | 26.0 |
| | Anaeromyces robustus v1.0<br>[jgi\|Anasp1\|325830] | GH48-DOC2-DOC2 | NaN | NaN | NaN | 22.7 |
| | Anaeromyces robustus v1.0<br>[jgi\|Anasp1\|293207]* | GH9-DOC2-DOC2<br>GH9-DOC2-DOC2-DOC2 | NaN | NaN | NaN | 21.6 |
| | Piromyces finnis v3.0<br>[jgi\|Pirfi3\|344383]* | DOC2-DOC2-GH6<br>GH6 | NaN | NaN | 23.3 | NaN |

| | | | | | | |
|---|---|---|---|---|---|---|
| *3* | Piromyces finnis v3.0 [jgi\|Pirfi3\|354732]* | **DOC2-DOC2-GH6**<br>**GH6** | 24.6 | 22.1 | 25.7 | 24.5 |
| | Piromyces finnis v3.0 [jgi\|Pirfi3\|413312]* | GH48-DOC2-DOC2 | NaN | NaN | 23.4 | NaN |
| | Piromyces sp. E2 v1.0 [jgi\|PirE2_1\|23096]* | CE3-GH11-CE3<br>GH11<br>GH11-GH11<br>GH11-GH11-DOC2<br>GH11-GH11-GH11 | NaN | NaN | NaN | 22.4 |
| | Piromyces sp. E2 v1.0 [jgi\|PirE2_1\|47957]* | DOC2-DOC2-GH43<br>GH43 | NaN | NaN | NaN | 23.5 |
| | Piromyces sp. E2 v1.0 [jgi\|PirE2_1\|11019]* | GH124-DOC2<br>GH124-DOC2-DOC2 | NaN | NaN | NaN | 22.0 |
| *4* | Piromyces sp. E2 v1.0 [jgi\|PirE2_1\|12703]* | **GH48-DOC2**<br>**GH48-DOC2-DOC2** | NaN | NaN | 25.7 | 25.9 |
| *5* | Piromyces sp. E2 v1.0 [jgi\|PirE2_1\|21620] | **GH8-DOC2-DOC2** | NaN | NaN | 25.7 | 25.8 |
| | Piromyces sp. E2 v1.0 [jgi\|PirE2_1\|20181]* | GH9-DOC2-DOC2-DOC2 | NaN | NaN | NaN | 24.1 |
| | Neocallimastix californiae G1 v1.0 [jgi\|Neosp1\|699681]* | CBM13-DOC2-DOC2-CE1<br>CE1<br>DOC2-CE1<br>DOC2-DOC2-CE1 | NaN | NaN | 23.6 | NaN |
| *6* | Neocallimastix californiae G1 v1.0 [jgi\|Neosp1\|177128]* | **DOC2-DOC2-DOC2-GH5_1**<br>**DOC2-GH5_1GH5_1** | 21.5 | NaN | 22.4 | 22.0 |
| | Neocallimastix californiae G1 v1.0 [jgi\|Neosp1\|514294]* | DOC2-DOC2-GH5_4-GH5_4-GH5_4-DOC2-DOC2<br>GH5_4-GH5_4-DOC2-DOC2 | NaN | 22.1 | NaN | NaN |
| | Neocallimastix californiae G1 v1.0 [jgi\|Neosp1\|708206]* | GH2<br>GH2-CBM13<br>GH2-CBM13-DOC2<br>GH2-CBM13-DOC2-DOC2 | NaN | NaN | 23.3 | NaN |
| | Neocallimastix californiae G1 v1.0 [jgi\|Neosp1\|699432] | GH3-DOC2-GH6-DOC2-DOC2 | NaN | 21.4 | 23.1 | NaN |
| | Neocallimastix californiae G1 v1.0 [jgi\|Neosp1\|503798] | GH45-DOC2-DOC2-DOC2 | NaN | NaN | NaN | 20.8 |
| | Neocallimastix californiae G1 v1.0 [jgi\|Neosp1\|704451] | GH48-DOC2-DOC2 | NaN | NaN | 24.4 | NaN |
| | Neocallimastix californiae G1 v1.0 [jgi\|Neosp1\|705896] | GH48-DOC2-DOC2 | NaN | NaN | 23.7 | NaN |
| | Neocallimastix californiae G1 v1.0 [jgi\|Neosp1\|675650]* | GH8-DOC2<br>GH8-DOC2-DOC2 | NaN | 25.0 | NaN | NaN |
| | Orpinomyces sp. [jgi\|Orpsp1_1\|1178620]* | DOC2-DOC2 | NaN | NaN | NaN | 22.2 |
| | Orpinomyces sp. [jgi\|Orpsp1_1\|1188096]* | DOC2-DOC2 | NaN | NaN | 22.5 | NaN |
| | Orpinomyces sp. [jgi\|Orpsp1_1\|1188535] | DOC2-DOC2 | NaN | NaN | NaN | 25.4 |
| | Orpinomyces sp. [jgi\|Orpsp1_1\|1176933]* | DOC2-DOC2-DOC2-GH6 | NaN | NaN | NaN | 23.1 |
| *7* | Orpinomyces sp. [jgi\|Orpsp1_1\|1179777]* | **DOC2-DOC2-DOC2-GH9**<br>**GH9** | 23.7 | 23.0 | NaN | NaN |
| *8* | Orpinomyces sp. [jgi\|Orpsp1_1\|1175496] | **DOC2-DOC2-GH6** | NaN | NaN | 24.2 | 23.6 |
| | Orpinomyces sp. [jgi\|Orpsp1_1\|1178177]* | DOC2-DOC2-GH6<br>DOC2-GH6 | NaN | NaN | 21.6 | NaN |
| | Orpinomyces sp. [jgi\|Orpsp1_1\|1176624]* | DOC2-DOC2-GH6<br>DOC2-GH6;GH6 | NaN | NaN | NaN | 20.9 |
| | Orpinomyces sp. [jgi\|Orpsp1_1\|1176522]* | GH3-DOC2-GH6-DOC2-DOC2 | NaN | NaN | NaN | 23.3 |
| | Orpinomyces sp. [jgi\|Orpsp1_1\|1182666]* | GH43<br>GH43-CBM6-CBM13<br>GH43-CBM6-CBM13-DOC2<br>GH43-CBM6-CBM13-DOC2-DOC2<br>GH43-CBM6-DOC2<br>GH43-CBM6-DOC2-DOC2 | NaN | NaN | NaN | 24.6 |

| | | | | | | |
|---|---|---|---|---|---|---|
| *9* | Orpinomyces sp. [jgi\|Orpsp1_1\|1182381] | **GH48-DOC2-DOC2** | NaN | 23.9 | 26.6 | 25.9 |

**Table S3:** Metatranscriptome Quality Filtering Stats.

| | **Total reads [count(%)]** | **Low quality reads [count(%)]** | **Artifact reads [count(%)]** | **rRNA reads [count(%)]** | **Remaining reads [count(%)]** |
|---|---|---|---|---|---|
| **Rumen Metatranscriptome** | 423,409,432 (100) | 3000746 (4.78) | 119751 (0.15) | 4245680 (5.26) | 379,212,367 (89.56) |

**Text S1:** FASTA sequences of the putative auxiliary proteins (three bacterial extracellular solute-binding proteins and one oxidoreductase), detected within the metaproteome. This list also includes one additional viral protein sequence detected at the upper range of the protein detection level.

Bacterial extracellular solute-binding proteins (3):

```
>Vir_gene_id_63857
GDQGTRAIINNLYGGTFTNPEHTAYTADSPENIKALELLQSLEGVYFNAAENGGEEITAFRQGLLKMAFCWNIAQ
QLPGETTGDRTYDDEEIVFMSFPAEDGVAALCGGIWGFGIFDNGDAAKIEAAKEFIKYFCDGAGTEAAVKTAKYF
AVRDVADGKDISGIWADDDTMNAYKVLMPYLGDYYQVTSGWATARTEWWNMLQRIGDGGDVATEVAVFCANANAA
AAA

>Vir_gene_id_63859
MLTLSExxxxLWTYPIGQWGNADAVGALMADFEAATGIKVTVEYLDYTNGDDQVNMAIEGKNAPDLVMEGPERLV
ANWGAKGYMVDLADLWDEEDLAQVNPSCVSACFTADGACYEYPV

>Vir_gene_id_63912
MKKVLALVLAMVMVLSLVSFASADDKVISIFLGGGTPLSMDPALNSASAGSNIIRSAFAGLTGFQYNEAGEPEMA
PEIAESYEVSEDGLTYSFVLRENLKWSDGTDCTASQIKASWERAASAELGADYGFLYDVISRAEDGSLAIDVDDA
ARTFVVHLPQPCAYFLDLCAFVPFYPVRVDLADNEGIWATNPETYVGLGAFKMTKYAVDDVISFEKNPYYWNADA
VKLSGLNFYLSEDNTAILTAYENGTVQYIQSISSSEFDRLNATYPGELAFWPTQGTYYILFNVHKDLSPASKQLT
VQEQSKARFALGELINRYEVVTYVTKGGEVAATGFFPAGLADGLNSDVRAAEGYGVWYTGTNEFSDVNPDYTVDQ
VEALQTLVDLGYPYTGSIEGGDIVFTDFPSIEFAFNNSGNNALIIQYVQETWNQFGITGVVNQEAWATLQSKLKA
GDAEAARMGWIADFNDVVNFLEIFISASGNNYPRLGREIGDYTRNSEVTADAGKGAYWGPNGDQTWAEAYDALVD
AVKAATDPVERASLAAEAEKVLMATGGVNPLYYYTTAQMLKPNVTDVIRLATGDVIWTYADIN
```

Oxidoreductase (1):

```
>Vir_gene_id_71822
IYALEHGIPVLLEKPFSVTLEEAEAVCRAEKASGKFVSVGFQPRFDANMQMIKKIVDSGVLGKIYYIQTGGGRRR
GIPGSTFIEKSTGGIGALGDIGCYSLDMVLNAIGYPKPLTVSGYISDFFGKNPKYNGKDAERFSVDDFAAAFIRL
EGDIILDFRIAWAMHVNTPGDTIIFGTEGALRIPSTDCxxxxRSRGSPTG
```

Putative porter protein (high protein detection level; $\log_2$(LFQ) score = 31.5) (1):

```
>Vir_gene_id_42007
MWSAIVQKIKEILQKMIGKNTLEQTLHVSPVISSEMENAIQLWGDMYKGNPPWVHEPTWEDPSRVVSLGIPALVA
SEKARTALLEFSSEVTTPIEEVEVQNPNYQPPMQDEFGVIKPEIGTPTVKESKPVGNTERADFLNENYKWLKKRL
RTQIEYGIAKGGLVIKPYLIQRKGEWAFEYDYIQADAFYPLSFDNNGDITEAAFLQTHKDKEFVYTRVEYHKWVD
NVVTVINRAFKTTANVHDTNGIDLGMEVPLTEVNEWKDLEPTTTITDVDRPLFAYFKMPEANTIDTTSSLGVSGY
SRAVQLIRDADEQYSRLLWEYEGGEMAIDVDRDALKFIEGSDGTVHTVLPKLQQRLFRKIDLGNDDVYNPFAPTL
RDAQYIQGLNAILMRIEDVTGLSRGTLSDVTIEAKTATELKMLKQRSYQTNEHIQEAIQTTLEDVVYIMNVYCDL
YDVTPDGEYEISFEWDDSLITDKDDELGKRITLQQNGLASRVENRMWYFGETERQAMEALAKIDEENAKQAMANM
ELEQSSQALHENNNGNTTGMNNETPNKNQPNKIKGVDKNNAQ
```
